# HD Spot: Interpretable Deep Learning Classification of Single Cell Transcript Data

**DOI:** 10.1101/822759

**Authors:** Eric Prince, Todd C. Hankinson

## Abstract

High throughput data is commonplace in biomedical research as seen with technologies such as single-cell RNA sequencing (scRNA-seq) and other Next Generation Sequencing technologies. As these techniques continue to be increasingly utilized it is critical to have analysis tools that can identify meaningful complex relationships between variables (i.e., in the case of scRNA-seq: genes) in a way such that human bias is absent. Moreover, it is equally paramount that both linear and non-linear (i.e., one-to-many) variable relationships be considered when contrasting datasets. HD Spot is a deep learning-based framework that generates an optimal interpretable classifier a given high-throughput dataset using a simple genetic algorithm as well as an autoencoder to classifier transfer learning approach. Using four unique publicly available scRNA-seq datasets with published ground truth, we demonstrate the robustness of HD Spot and the ability to identify ontologically accurate gene lists for a given data subset. HD Spot serves as a bioinformatic tool to allow novice and advanced analysts to gain complex insight into their respective datasets enabling novel hypotheses development.

## MAIN

High throughput data has become increasingly ubiquitous in biomedical research, with advances in computer science facilitating the ability of biomedical scientists to complete analyses. Vast clinical datasets are routinely generated in fields such as radiology, and basic science datasets like those yielded by Next Generation Sequencing (NGS) technologies (i.e., single-cell RNA-sequencing [scRNA-seq]) have become relevant in clinical care. In the case of NGS data, current analysis methodologies suffer from a lack of standardization. While some genetic signatures are easy to interpret due to their linear nature, a far greater proportion of genetic interactions are non-linear and therefore not reliably interpretable using manual techniques. Further, novel bioinformatic findings increasingly require the concatenation of multiple high-level mathematical techniques in order to filter out a manageable subset of features of interest. Current methods suffer from both human bias and sampling error present within the dataset. These human biases can arise from an analyst not being blind to the dataset and therefore preferential to a desired outcome, or potentially applying mathematically incorrect/sub-optimal models to the target dataset.

Deep Learning (DL) is a subset of machine learning that is capable of concurrently modeling simple and complex mathematical relationships present within a given dataset. DL – and machine learning in general – is being applied across myriad scientific disciplines with success levels often superior to those of human experts. The DL landscape is vast, although certain architectures are becoming established for given problem sets (e.g., Convolutional Neural Networks [CNN] for image classification and object detection, Long Short-Term Memory [LSTM] networks for sequence-based analysis, and autoencoder networks for anomaly detection). Though methodology and architecture standards of practice are emerging, the successful implementation of these DL models can be elusive.

Among the challenges in developing a reliable DL system is optimization of the various meta-parameters. These parameters include, but are not limited to, learning rate scale and behavior, dropout and regularization, optimizers, and activation functions. Selection of the optimal settings of such parameters is made more complex by the fact the parameters are usually continuous, and their relationships are often non-linear. In order to address this challenge, meta-optimization solutions that can intelligently explore the parameter optimization space have been developed. A promising approach is modeled after the simple Genetic Algorithm (GA)^1^. By treating each possible parameter set as an “individual” and assigning a “fitness” value (i.e. accuracy in a classification task), one can test a population of individuals, rank them by their fitness, and then create a new population using the parameters found in the fittest individuals. As the process iterates for several generations, the most important parameter selections become homogenous and less important parameters become more heterogenous. This approach is significantly less resource intensive than a comparable brute-force approach, which examines all possible parameter value combinations.

We present a computational method, termed HD Spot, that leverages DL architectures and GA meta-optimization to create optimized novel classifiers and generate feature importance information for high-throughput (HT) datasets. HD Spot provides an easy-to-use framework for scientists to interrogate differences in HT data (e.g., bulk RNA-seq transcript differences between disease subtypes, identifying interesting genes in unknown scRNA-seq cell clusters with non-canonical signatures, etc.). Further, HD Spot helps to remove the potential human bias that can be present in advanced analytical pipelines. Finally, by combining HD Spot results with established techniques, users can identify potential dynamic relationships within a dataset, thereby improving the value of their analysis.

## RESULTS

### Binary scRNA-seq Dataset Performance

To evaluate the robustness of HD Spot in a binary classification scenario, we investigated an integrated dataset containing two different sets of control versus IFN-β stimulated immune cells (CD14+ Monocytes and CD4+ Naïve T cells [Figure 1a]). The gene expression signatures of these two populations are not easily differentiable, even when transformed using the non-linear dimension reduction technique UMAP (Figure 1b). The maximum mean performance (AUPR) of HD Spot, when analyzed using 5-fold cross-validation, was 0.587 and 0.881 for CD4+ T cells and CD14 Monocytes, respectively. To estimate the ontology differences between control and stimulated populations of CD4+ T cells, the top-20 most important (i.e., highest SHAP values; Figure 1c) genes were submitted for Metascape analysis. The resultant list of significant ontologies suggests that the Naïve CD4+ T cell (T_h0_) population is being activated to the T_h2_ phenotype via IL-4 (Figure 1c; CD4+ Memory T Cells, yellow highlight). Similarly, we collected the top-20 most important genes that distinguish control from IFN-β stimulated CD14 Monocytes and submitted these for Metascape analysis (Figure 1c). Significant ontologies suggest the population contains Monocytes that are being activated (Figure 1c; CD14 Monocytes, yellow highlight). By utilizing SHAP values, we are also able to investigate variable interactions that help differentiate the populations of interest. For example, in CD4+ T cells we can see that increase in TNFSF10 combined with a decrease in RSAD2 expression suggests the cell is stimulated, while the absence of TNFSF10 indicates that the cell belongs to the control group (Figure 1d). However, these interactions can still be difficult to interpret such as the relationship between CCL2 and DHRS9 in CD14 Monocytes (Figure 1e). While it is known that DHRS9 and CCL2 play a role in CD14 Monocyte activation^2^, the interaction between expression levels in these genes is convoluted.

**Figure 1.**
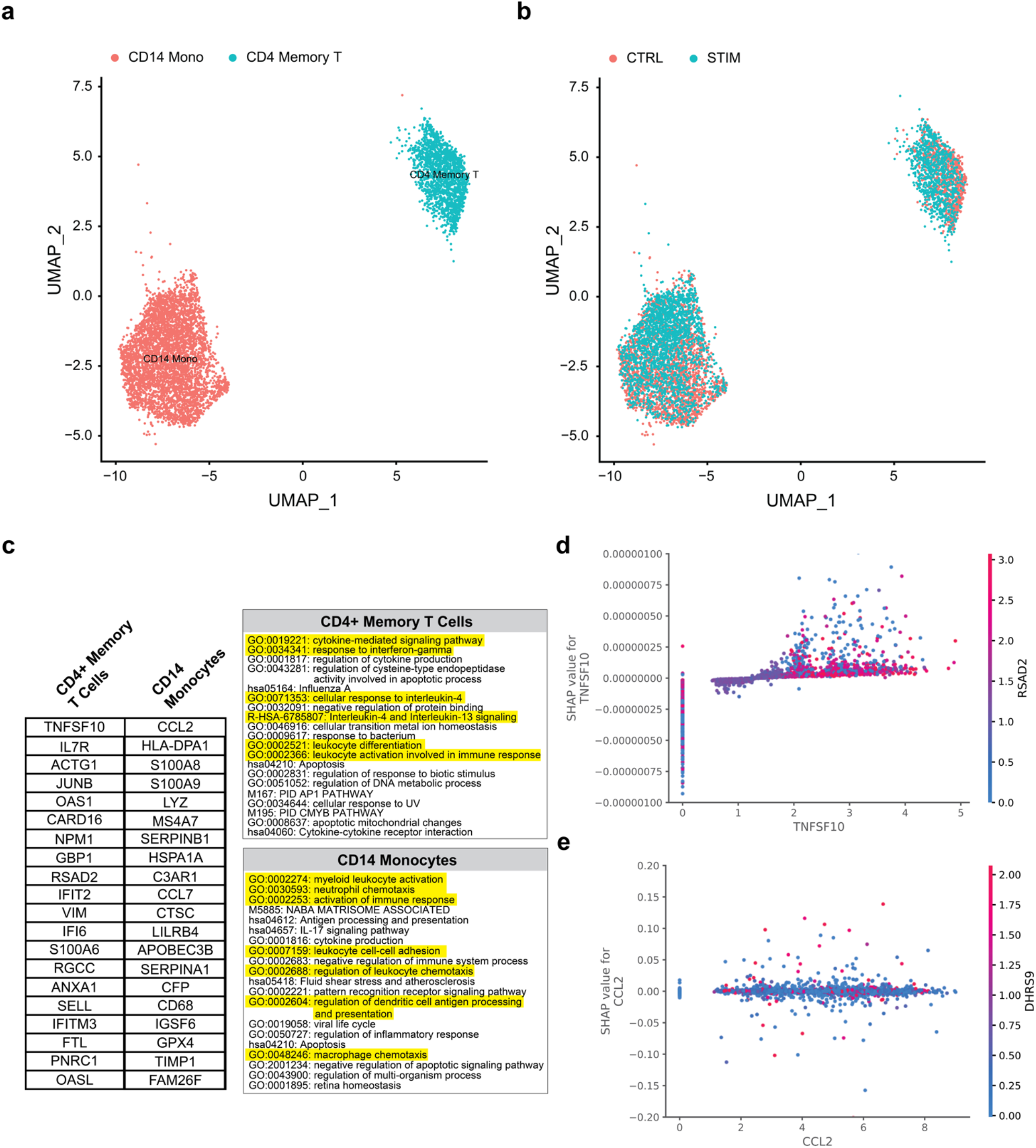
Classification and Feature Importance of IFN-β Treated PBMCs. **a.** UMAP of CD4+ memory T cells (blue) and CD14 monocytes (red). **b.** UMAP of non-treated (CTRL; red) and IFN-β treated (STIM; blue) CD4+ memory T cells and CD14 monocytes. **c.** Top-20 most important (largest SHAP value) genes as determined by HD Spot for CD4+ memory T and CD14 Monocytes, and respective ontology enrichments as determined by Metascape. **d.** Interaction plot for CD4+ Memory T Cell data regarding TNFSF10 and RSAD2. SHAP values > 0 indicate that the interaction is related to the STIM genetic profile, SHAP values < 0 indicate the inverse. **e.** Interaction plot for CD14 Monocyte data regarding CCL2 and DHRS9. SHAP values > 0 indicate that the interaction is related to the STIM genetic profile, SHAP values < 0 indicate the inverse. TNFSF10 and CCL2 maintain top-ranked SHAP values for CD4+ Memory T and CD14 Monocyte cells, respectively. RSAD2 and DHRS9 were estimated to have the greatest SHAP interaction values with TNFSF10 and CCL2, respectively.

### Multiclass Performance on scRNA-seq Dataset

Using the raw feature counts from the 10X 2700 Peripheral Blood Mononuclear Cell (PBMC) scRNA-seq dataset as labeled in the Seurat Vignette (Figure S1), HD Spot generated a multiclass classifier for the nine different cell types. For each cell type, the top twenty most significant genes utilized in the classification process was determined (Figure 2a). For each top twenty gene list, a Metascape analysis was performed (Figure 2b-c). HD Spot independently identified genes that were referenced as cell type markers in the Seurat Vignette. Specifically, it identified S100A4 in CD4+ memory T cells, MS4A1 in B cells, LYZ in CD14+ Monocytes, GNLY and NKG7 in Natural Killer (NK) cells, and FCGR3A in FCGR3A+ Monocytes. Moreover, Metascape analysis indicated expected phenotypic qualities. For example, *cytokine-mediated signaling* pathway enrichment in CD4+ memory T cells, *Antigen processing and presentation* in B cells, *positive regulation of apoptotic process* in NK cells, *lymphocyte apoptotic process* in CD8+ T cells, and *phagocytosis* and *response to interferon-gamma* in FCGR3A+ Monocytes. Classes that were highly imbalanced (dendritic cells [n=32] and platelets [n=14], due to the relatively low frequency that they occur in the dataset) were not correctly classified using raw feature counts as input values. The top-20 genes in these scenarios were similar to the most variant genes as identified by PCA (Figure S2). Interestingly, in the multiclass scenario HD Spot identified many of the same genes identified as important in Principle Component 1 (i.e., the variables with the largest eigenvalues in the first dimension of Principle Component Analysis [PCA]; Figure S2). This reflects model precision, as autoencoders that optimally learn a data manifold (i.e.,landscape) decompose data similarly to PCA^3^.

**Figure 2.**
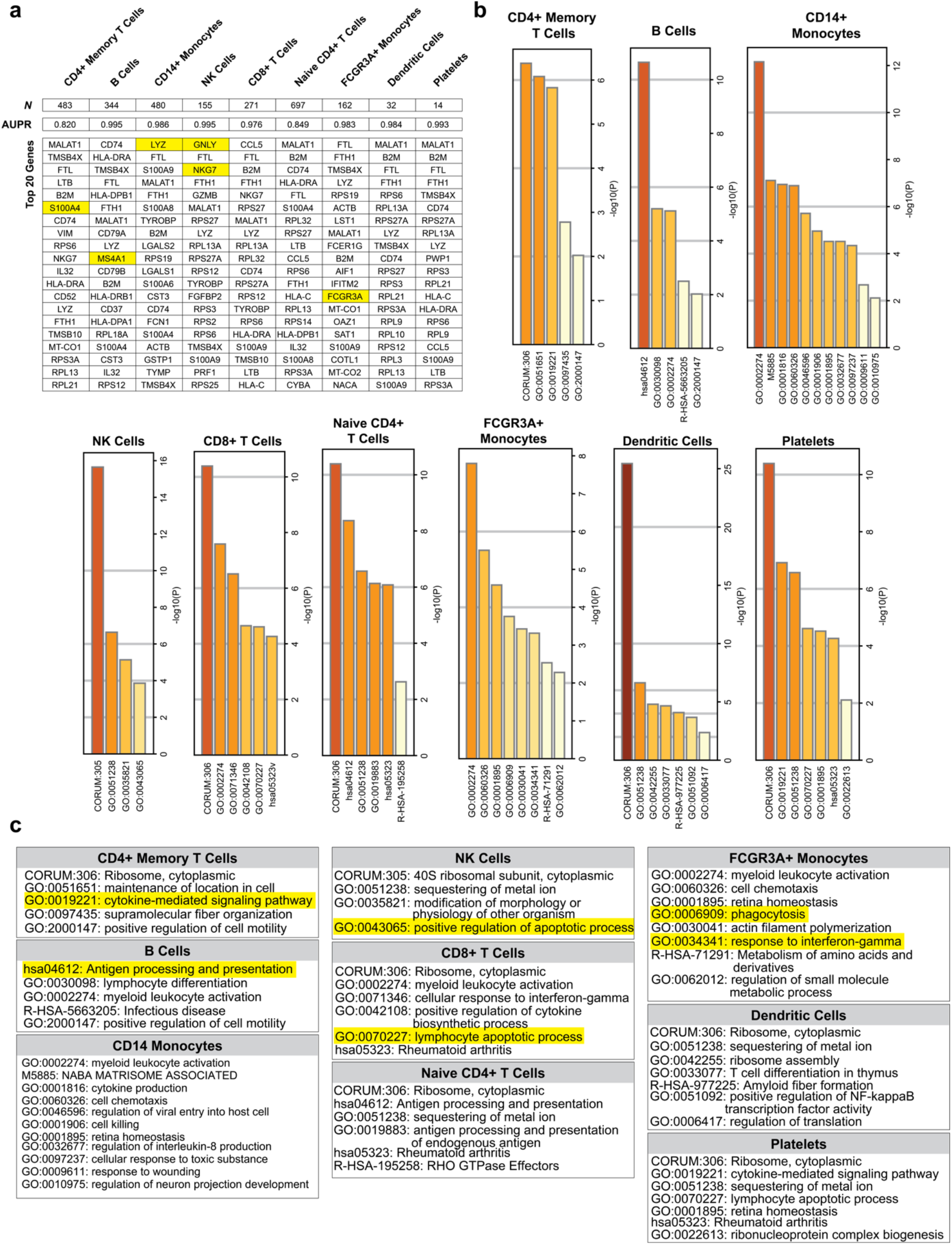
HD Spot Analysis of 10X 2700 PBMC Dataset Raw Feature Counts. **a.** Top-20 most important (largest SHAP values) transcripts for each class as determined in a one-versus-all approach for each class (i.e., cell type). **b.** Metascape-determined significant ontologies based on HD Spot-identified gene lists. **c.** Long-form names of GO, KEGG, and Reactome pathways returned in Metascape analysis. Values that are highlighted yellow correspond with genes and ontologies discussed in the Results section.

### Comparison of Single Cell Feature Counts and Single Cell Scaled Expression Values

The 10X 2700 PBMC dataset was normalized and submitted for HD Spot analysis. The 20 most significant genes for each cell type (Figure 3a) and the significantly enriched ontologies (Figure 3b-c) were identified. While raw feature counts generated 62 significant ontologies across the nine cell types, only 39 (63%) were unique. With normalized expression values, the sequence of HD Spot followed by Metascape analysis returned 50 significant ontologies, of which 48 (96%) were unique. This suggests that the quality of information is enhanced when submitting normalized data to HD Spot. Evidence for quality of information improvement is also observed in the ontologies and top 20 genes for each cell type. For example, using raw feature counts the CD4+ memory T cell phenotype is suggested through enrichment of cytokine-mediated signaling *(GO:0019221).* Using normalized expression values, the same pathway is enriched, but other pathways that have known associations with CD4+ memory T cells are also identified. These include *Tumor Necrosis Factor-mediated signaling pathway*^4^, *Epstein-Bar virus infection*^5^, and *lymphocyte differentiation*^6^. This improved quality of information can also be seen in B cells^7^, in CD14 Monocytes, in NK cells^8^, in CD8+ T cells, in CD4+ naïve T cells^9^, and in FCGR3A+ Monocytes. While HD Spot failed to yield accurate estimates for dendritic cells and platelets when using raw feature counts, it returned more phenotypically consistent genes lists when using scaled expression values. For example, Dendritic cells have known interactions with translation initiation complex formation^10^ and are activated in response radiation^11^, both of which were identified. Additionally, CCDC94 (YJU2) also has relatively high expression in dendritic cells according to BioGPS^12^. Finally, platelets are known to be activated via reactive oxygen species and ferroptosis^13^ and play a role in angiogenesis^14^.

**Figure 3.**
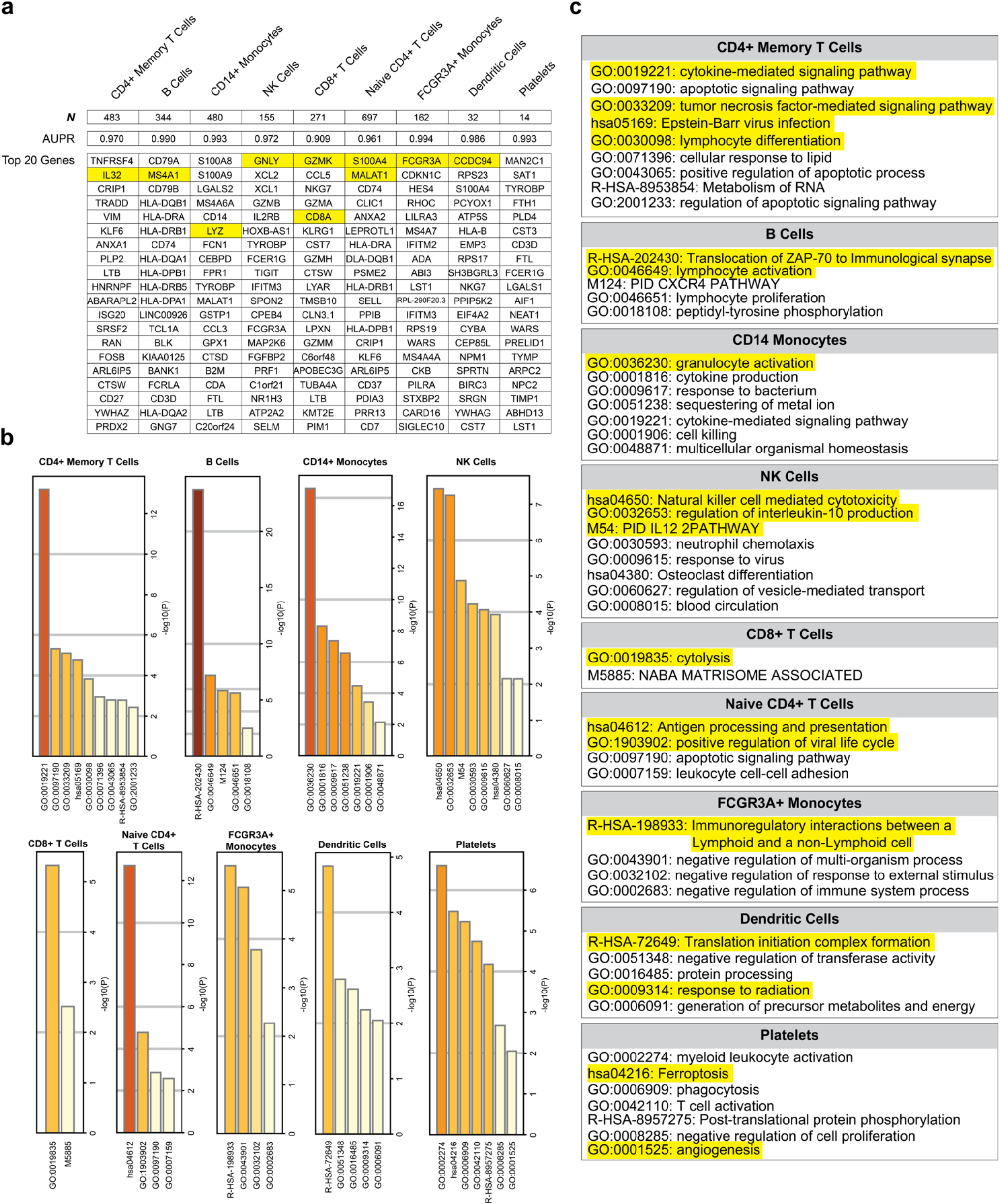
HD Spot Analysis of 10X 2700 PBMC Dataset Normalized Expression Values. **a.** Top-20 most important (largest SHAP values) transcripts for each class as determined in a one-versus-all approach for each class (i.e., cell type). **b.** Metascape-determined significant ontologies based on HD Spot-identified gene lists. **c.** Long-form names of GO, KEGG, and Reactome pathways returned in Metascape analysis. Values that are highlighted yellow correspond with genes and ontologies discussed in the Results section.

## Discussion

This manuscript preliminarily assesses a deep learning approach for analyzing complex linear and non-linear relationships in high-dimensional data, such as scRNA-seq, in a manner that is free of human bias. The impetus was to create a method, downstream of clustering, to distinguish cell populations that lack canonical gene signatures, and may be difficult to separate. Using two scRNA-seq datasets, we demonstrated that HD Spot inferred meaningful gene importance rankings, which estimated the given scenarios well. In the context of feature counts versus scaled data, both scenarios yielded genes and enriched ontologies that accurately infer at least some of the cell types. Scaled data resulted in improved classification performance, enhanced quality of information, and was more robust in the classes with low sample sizes (dendritic cells and platelets).

In addition to statistically describing unknown cell populations (data classes), there are other applications for this approach. HD Spot lowers the barrier for interpretation of complex data by enabling researchers who lack sophisticated computational experience to analyze their own dataset without the need for a bioinformatician. HD Spot is also well suited as an adjunct for advanced bioinformaticians. For example, it can be used to generate statistical insight regarding data relationships, to validate analytical findings, or to generate new potential research directions. Additionally, Linneaus can serve as an optimized classifier that can compare and contrasting multiple datasets. For example, a previously analyzed dataset can be directly compared to a new, non-analyzed, dataset of the same type and variable structure, in order to determine how the datasets are related.

An obvious limitation of HD Spot is the computational demand. The algorithm was developed using high performance computing (HPC)-caliber hardware, with multiple accessible GPUs. Real-time turnaround is reduced by asynchronously evaluating models within each Genetic Algorithm iteration. A potential method to decrease this computational requirement would be to implement an early stopping method to the GA. In this case, the GA would only proceed if model fitness was improving, thus eliminating potentially unnecessary training. Another limitation in our approach is that we have characterized immune cell types that are more easily differentiated than cell types in many other biological contexts. For example, subtyping within a cell or tumor type may prove to be a greater challenge than the proof of principle we present here. Future work to address this would be to stress test the HD Spot algorithm across more complex scRNA-seq datasets, as well as exploring other types of high-throughput biomedical data (i.e., ATAC-seq, high-dimension flow cytometry, multi-color histology, etc.).

## Methods

### Summary

HD Spot (of which the “Spot the Difference” picture game is the eponym) is a three-step machine learning process that seeks to optimize a classifier for a given dataset and then determine which variables are most important in the decision-making process (Figure M1). All models were constructed using the TensorFlow 2.0 (TF2) python framework and are cross-compatible with CPU, GPU, and/or TPU-based systems. First, using a simple Genetic Algorithm (GA) as a meta-optimization framework, an autoencoder is optimized to a custom fitness function (details below) in order to compress data information for classification. Second, weights from the encoder component of the optimized autoencoder are transferred to a classifier network, which is subsequently optimized via GA. Third, using the python Shapley Additive Explanations (SHAP) module, dataset feature importance is evaluated as Shapley values^15^.

**Figure M1.**
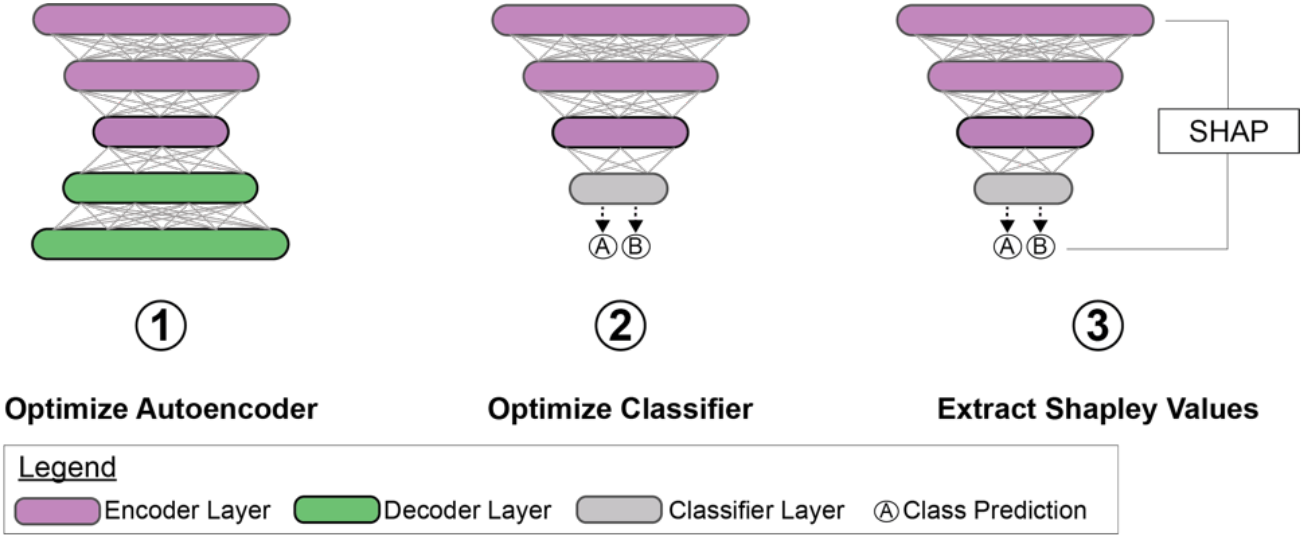
High-Level Overview of HD Spot Framework. Weights from optimized encoder layers are used to initialize classifiers which are then optimized. Feature importance is determined using Shapley values via the python SHAP module.

### Data Acquisition

Validation of HD Spot efficacy was achieved utilizing two datasets published with the R-based Seurat single-cell analysis framework^16,17^. The first dataset (control versus IFN-β-stimulated PBMCs) was generated by integrating two single cell RNAseq datasets^18^ as described in the Seurat vignette^19^ using the SeuratData R package. The second dataset was sourced from 10X genomics and it is known as the 2,700 Peripheral Blood Mononuclear Cells (PBMCs) dataset. The Seurat object for this dataset contains feature counts and scaled data (scaled via the Seurat ‘LogNormalize’ function) with designated cell type labels and is publicly available^20^. For each dataset, values were extracted and transposed such that columns are genes and rows are Unique Molecular Identifiers (UMIs; i.e., cells). The right-most column is set to contain the class label (i.e., cell type) as given in the published datasets. Finally, datasets were exported as comma-delimited tables.

### Genetic Algorithm Meta-optimizer

The GA was adapted from a publicly available repository^21,22^. The general algorithm is as follows:

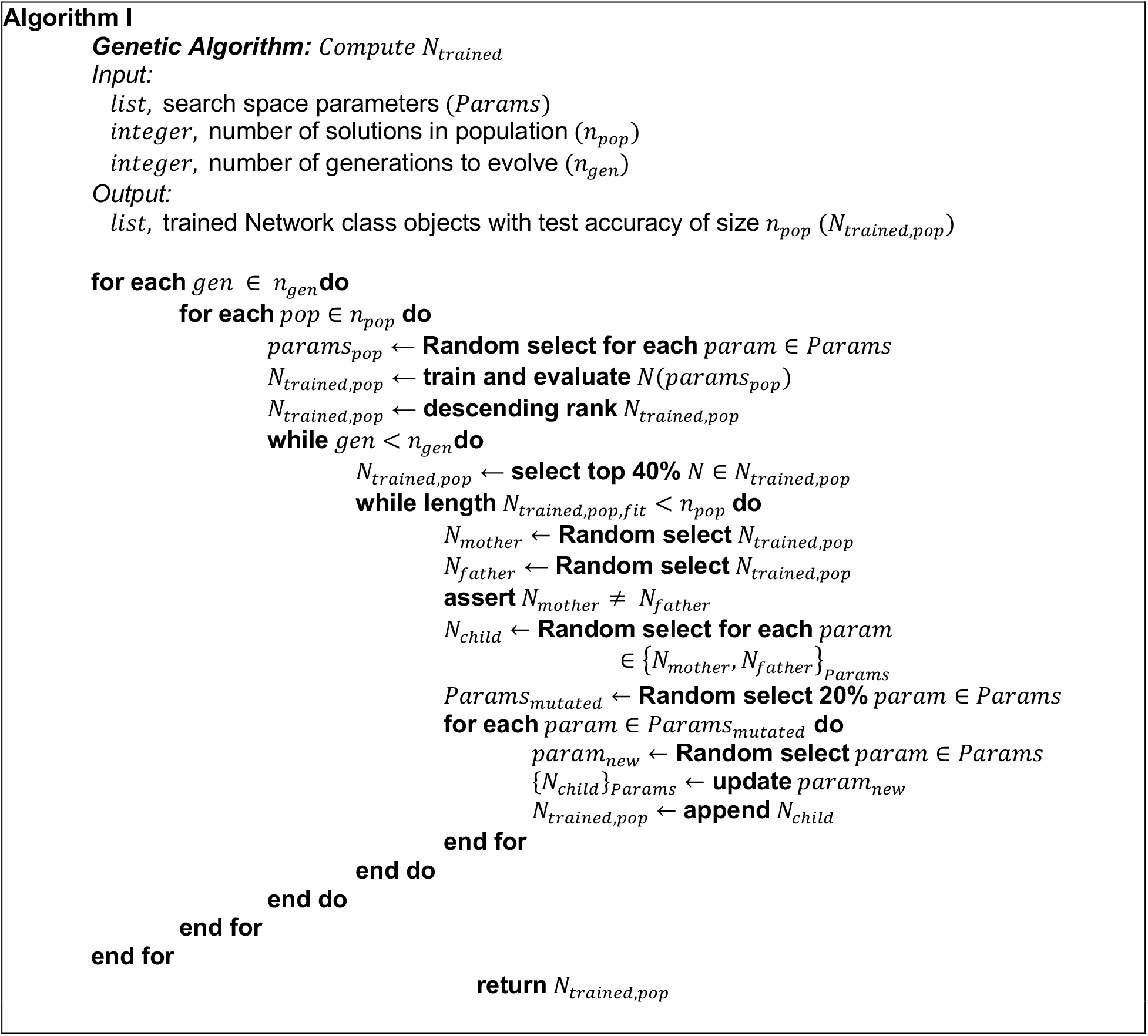

### Defining the GA Optimization Search Space

By default, the GA utilizes a pool of variables that include architectural and learning algorithm features. This parameter pool is slightly different for the autoencoder (Table 1) and the classifier networks. Specifically, classifier networks are given two additional parameters that are a selection of the top 5 optimized autoencoders and the ability to continue to train the encoder layer in the classifier or just extract the feature embeddings (i.e., trainable or not, respectively). The distinction between *inner* activation and *final* activation is that final activation is the activation of the final layer, and inner is the activation of all other previous layers. Additionally, all models were trained for 1,000 epochs but with early stopping, which was triggered either when values were incomputable (*NaN*) or when 5 epochs occurred without model performance improvement. This allowed us to ignore the training duration variable during model optimization.

**Table 1.** Parameter Pool for GA Optimization

| Batch Size | Learning Rate | Dropout Rate | Inner Activation | Final Activation | Optimizer | L1/L2-Kernel/Bias/Activity Weight |
| --- | --- | --- | --- | --- | --- | --- |
| 128 | 0.1 | 0.0 | Sigmoid | Sigmoid | RMSProp | 0.0 |
| 256 | 0.01 | 0.2 | Softplus | Softplus | AdaGrad | 0.1 |
| 512 | 0.001 | 0.5 | Softsign | Softsign | Adam | 0.01 |
| 1024 | 0.0001 |  | Softmax | Softmax | AdaDelta | 0.001 |
|  |  |  | EIU | EIU | AdaMax | 0.0001 |
|  |  |  | Hard Sigmoid | Hard Sigmoid | SGD | 0.00001 |
|  |  |  | Linear | Linear |  |  |
|  |  |  | SeLU | SeLU |  |  |
|  |  |  | Tanh | Tanh |  |  |

The decision to include the types of parameters utilized in HD Spot was driven by a desire to reinforce the model in small dataset scenarios (specifically dropout and regularization). Moreover, model fitness was determined using 5-fold cross-validation for the classifier. HD Spot, by default, monitors several metrics, including categorical accuracy, AUROC, AUPR, precision, recall, and top-*k* categorical accuracy (in multiclass scenarios). It utilizes AUPR as the fitness metric. This is because AUPR is robust in situations that are highly class imbalanced, but it is also inherently connected to AUROC and therefore is also effective in class balanced situations.

By default, the GA is set to perform 10 generations of populations containing 100 solutions. This means the GA will assess 1,000 solutions, which equates to 7.38×10^-6^% of the possible parameter combinations in the autoencoder optimization (*N* = 13,548,902,400) and 7.38×10^-8^% of the possible parameter combinations in the classifier optimization (*N* = 1,354,890,240,000). While omitting the majority of solutions, the selection of 100 solutions per population enables a minimum parameter frequency of 12.5 per population, meaning the population contains more than 10% of each parameter. The intuition is that parameter selections with more profound impact on model performance will become homogenous in later generations, and less important parameters will remain heterogeneous in top solutions. HD Spot computational time is decreased by performing iterative training/evaluation cycles in an asynchronous fashion using the python ray and asyncio modules. This is important due to TF2 not releasing memory when iterations are not handled in unique process IDs (PIDs), which will cause an out-of-memory error if not accounted for.

### Defining the Autoencoder Fitness Function

Autoencoders are utilized to compress the data information to a lower dimensionality, which enables the utilization of a smaller classifier. This is because more compressed data will require fewer, but more relevant, DL network nodes to learn the dataset objective function, when compared to a dataset with a high degree of sparsity. In HD Spot, autoencoders are utilized in an unsupervised manner, which means that the network is modeling the dataset in its entirety, and *not* considering class labels in the process. The loss function utilized is the Mean Squared Error (MSE) between input and output values, and there is no accuracy metric implemented. To employ the GA, we required a static value to rank-order, as the fitness function and the final loss is inadequate for this purpose. In the fitness function, we sought to derive a function that would reflect the qualitative characteristics of an ideal loss curve: a quick training duration (i.e., rapid convergence of the objective function), and a low final loss. Intuitively, these characteristics reflect a model that has learned the data distribution in a stable manner. As such, the following fitness function was derived.

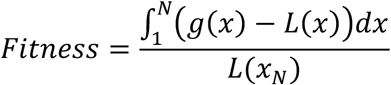

Where *x* is the number of epochs (*x*: {1,2,…, *N*}), *L*(*x*) is the loss function of the model over the training duration, *L*(*x_N_*) is the final loss value of training, and *g*(*x*) is a first-order polynomial (linear) regression mapped from *L*(*x*_1_) to *L*(*x_N_*). Importantly, if the slope of *g*(*x*) > 0, a penalty term (−1×10^9^) is multiplied with the fitness function to downrank such loss curves that are increasing. This function therefore transforms a series of metrics into a static method that will increase when a loss function has greater concavity (i.e., the fitness function has a larger numerator) and finishes with a lower final loss (i.e., smaller denominator).

### Optimizing Autoencoder

It is well known that autoencoders with three or more layers are difficult to successfully train. To overcome this, layers can be individually pretrained in the form of a Restricted Boltzmann Machine (RBM), which is an energy-based Markov Random Field model. In HD Spot, a new model is first created by pretraining each layer as an RBM using the contrastive-divergence method for 50 epochs. This number is arbitrary, though this is simply a layer weight initialization step and does not appear to require any tuning. Once model weights are initialized, the GA process is completed for the autoencoder, using the previously defined custom fitness function. Upon completion of GA optimization, the top 5 most-fit autoencoders are exported as saved models for downstream consideration.

### Optimizing Classifier

Classification networks were constructed using the encoder layers from the autoencoder as a foundation. Batch normalization, 2D convolution, 2D max pool, and a final softmax fully-connected layer were then appended to the encoder to create the final classifier architecture. Batch normalization was performed to improve model performance in small dataset or class-imbalanced scenarios. Convolution and max pool layers were utilized to extract more complex latent data relationships. Classifier optimization was then performed using the GA, as described above.

### Feature Importance Evaluation Using Shapley Values

The purpose of this effort was to develop an unbiased analytical method for high-throughput (i.e., high-dimensional) data capable of separating data into predetermined groups and subsequently returning the most important features (i.e., genes) the classifier used to separate the groups. To do so we utilize the python SHAP module to calculate Shapley additive values. This approach was selected due to the direct compatibility with TF2 models, as well as the ability to generate feature importance rankings and force plots to glean a linear insight into *how* features were utilized in the decision-making process.

### Metascape Analysis of Shapley Values

To determine enriched ontologies of genes filtered by importance, we utilized the Metascape online analysis tool^23^. Gene lists were submitted and analyzed as Homo sapiens in ‘Express Analysis’ mode.

### Computational Hardware

HD Spot was developed on an HPC server running a 10-core Intel Skylake Processor and 6 GPUs (4 NVIDIA Tesla K80 and 2 NVIDIA Tesla T4 GPU cards). On top of a CentOS 7 environment, we developed HD Spot using the publicly available TF2 docker image (https://hub.docker.com/r/tensorflow/tensorflow/).

### Code Availability

HD Spot is provided as-is without warranty on GitHub to encourage improvement and adaptation by the broader scientific community.

## ACKNOWLEDGMENTS

The authors wish to express their sincere gratitude towards The Morgan Adams Foundation Pediatric Brain Tumor Research Program for their donations that made the computational infrastructure available to develop this project. Author’s Note: The presented computational framework was initially titled “Linnaeus”. The authors wish to express their gratitude to Dr. Philipp Junker for illuminating the naming overlap with their scRNA-seq lineage tracing strategy^23^.

## Notes

#### Summary of Updates

v1.1: grammatical and formatting errors corrected. line numbers removed. v1.2: Linnaeus name-changed to HD Spot

## REFERENCES

1. Wynter, A. (2019). On the Bounds of Function Approximations https://dx.doi.org/10.1007/978-3-030-30487-4_32

2. Atri, C., Guerfali, F., Laouini, D. (2018). Role of Human Macrophage Polarization in Inflammation during Infectious Diseases International Journal of Molecular Sciences 19(6), 1801. https://dx.doi.org/10.3390/ijms19061801

3. Goodfellow, I., Bengio, Y., Courville, A. (2016). Deep Learning

4. Croft, M. (2009). The role of TNF superfamily members in T-cell function and diseases Nature Reviews Immunology 9(4), 271–285. https://dx.doi.org/10.1038/nri2526

5. Amyes, E., Hatton, C., Montamat-Sicotte, D., Gudgeon, N., Rickinson, A., McMichael, A., Callan, M. (2003). Characterization of the CD4+ T Cell Response to Epstein-Barr Virus during Primary and Persistent Infection The Journal of Experimental Medicine 198(6), 903–911. https://dx.doi.org/10.1084/jem.20022058

6. Gasper, D., Tejera, M., Suresh, M. (2014). CD4 T-Cell Memory Generation and Maintenance Critical Reviews in Immunology 34(2), 121–146. https://dx.doi.org/10.1615/critrevimmunol.2014010373

7. Gobessi, S., Laurenti, L., Longo, P., Sica, S., Leone, G., Efremov, D. (2006). ZAP-70 enhances B-cell-receptor signaling despite absent or inefficient tyrosine kinase activation in chronic lymphocytic leukemia and lymphoma B cellsBlood 109()

8. Mehrotra, P., Donnelly, R., Wong, S., Kanegane, H., Germew, A., Mostowski, H., Furuke, K., Siegel, J., Bloom, E. (1998). Production of IL-10 by human natural killer cells stimulated by IL-2 and/or IL-12 Journal of Immunology

9. Douek, D., Brenchley, J., Betts, M., Ambrozak, D., Hill, B., Okamoto, Y., Casazza, J., Kuruppu, J., Kunstman, K., Wolinsky, S., Grossman, Z., Dybul, M., Oxenius, A., Price, D., Connors, M., Koup, R. (2002). HIV preferentially infects HIV-specific CD4+ T cells Nature 417(6884), 95. https://dx.doi.org/10.1038/417095a

10. Lelouard, H., Schmidt, E., Camosseto, V., Clavarino, G., Ceppi, M., Hsu, H., Pierre, P. (2007). Regulation of translation is required for dendritic cell function and survival during activation The Journal of Cell Biology 179(7), 1427–1439. https://dx.doi.org/10.1083/jcb.200707166

11. Persa, E., Szatmári, T., Sáfrány, G., Lumniczky, K. (2018). In Vivo Irradiation of Mice Induces Activation of Dendritic Cells International Journal of Molecular Sciences 19(8), 2391. https://dx.doi.org/10.3390/ijms19082391

12. Wu, C., Orozco, C., Boyer, J., Leglise, M., Goodale, J., Batalov, S., Hodge, C., Haase, J., Janes, J., Huss, J., Su, A. (2009). BioGPS: an extensible and customizable portal for querying and organizing gene annotation resources Genome Biology 10(11), R130. https://dx.doi.org/10.1186/gb-2009-10-11-r130

13. NaveenKumar, S., SharathBabu, B., Hemshekhar, M., Kemparaju, K., Girish, K., Mugesh, G. (2018). The Role of Reactive Oxygen Species and Ferroptosis in Heme-Mediated Activation of Human Platelets ACS Chemical Biology 13(8), 1996–2002. https://dx.doi.org/10.1021/acschembio.8b00458

14. Patzelt, J., Langer, H. (2012). Platelets in Angiogenesis Current Vascular Pharmacology

15. Lundberg, S., Lee, S. (2017). A Unified Approach to Interpreting Model Predictions

16. Butler, A., Hoffman, P., Smibert, P., Papalexi, E., Satija, R. (2018). Integrating single-cell transcriptomic data across different conditions, technologies, and species Nature Biotechnology 36(5),411. https://dx.doi.org/10.1038/nbt.4096

17. Stuart, T., Butler, A., Hoffman, P., Hafemeister, C., Papalexi, E., Mauck, W., Hao, Y., Stoeckius, M., Smibert, P., Satija, R. (2019). Comprehensive Integration of Single-Cell Data Cell 177(7), 1888–1902. e21. https://dx.doi.org/10.1016/j.cell.2019.05.031

18. Kang, H., Subramaniam, M., Targ, S., Nguyen, M., Maliskova, L., McCarthy, E., Wan, E., Wong, S., Byrnes, L., Lanata, C., Gate, R., Mostafavi, S., Marson, A., Zaitlen, N., Criswell, L., Ye, C. (2017). Multiplexed droplet single-cell RNA-sequencing using natural genetic variation Nature Biotechnology 36(1), 89. https://dx.doi.org/10.1038/nbt.4042/

19. Satija, R. (2019). Tutorial: Integrating stimulated vs. control PBMC datasets to learn cell-type specific responses

20. Satija, R. 2,700 PBMC Seurat Final Object https://www.dropbox.com/s/63gnlw45jf7cje8/pbmc3k_final.rds?dl=0

21. Harvey, M. Neural Network Genetic Algorithm. https://blog.coast.ai/lets-evolve-a-neural-network-with-a-genetic-algorithm-code-included-8809bece164 Larson, W. Genetic Algorithms: Cool Name and Damn Simple. https://lethain.com/genetic-algorithms-cool-name-damn-simple/

22. Zhou, Y., Zhou, B., Pache, L., Chang, M., Khodabakhshi, A., Tanaseichuk, O., Benner, C., Chanda, S. (2019). Metascape provides a biologist-oriented resource for the analysis of systems-level datasets Nature Communications 10(1), 1523. https://dx.doi.org/10.1038/s41467-019-09234-6

23. Spanjaard, B., Hu, B., Mitic, N., Olivares-Chauvet, P., Janjuha, S., Ninov, N., Junker, J. (2018). Simultaneous lineage tracing and cell-type identification using CRISPR–Cas9-induced genetic scars Nature Biotechnology 36(5), 469–473. https://dx.doi.org/10.1038/nbt.4124

