## Supplementary figures and images for "HD Spot: Interpretable Deep Learning Classification of Single Cell Transcript Data"

### FigS1

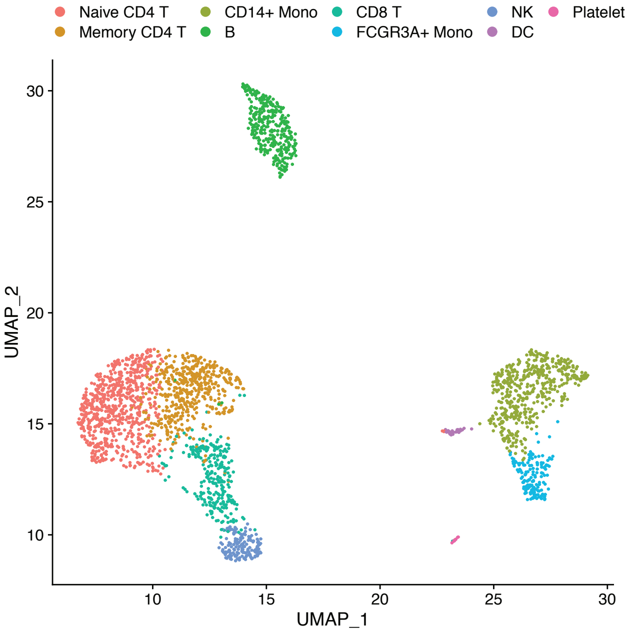

### FigS2

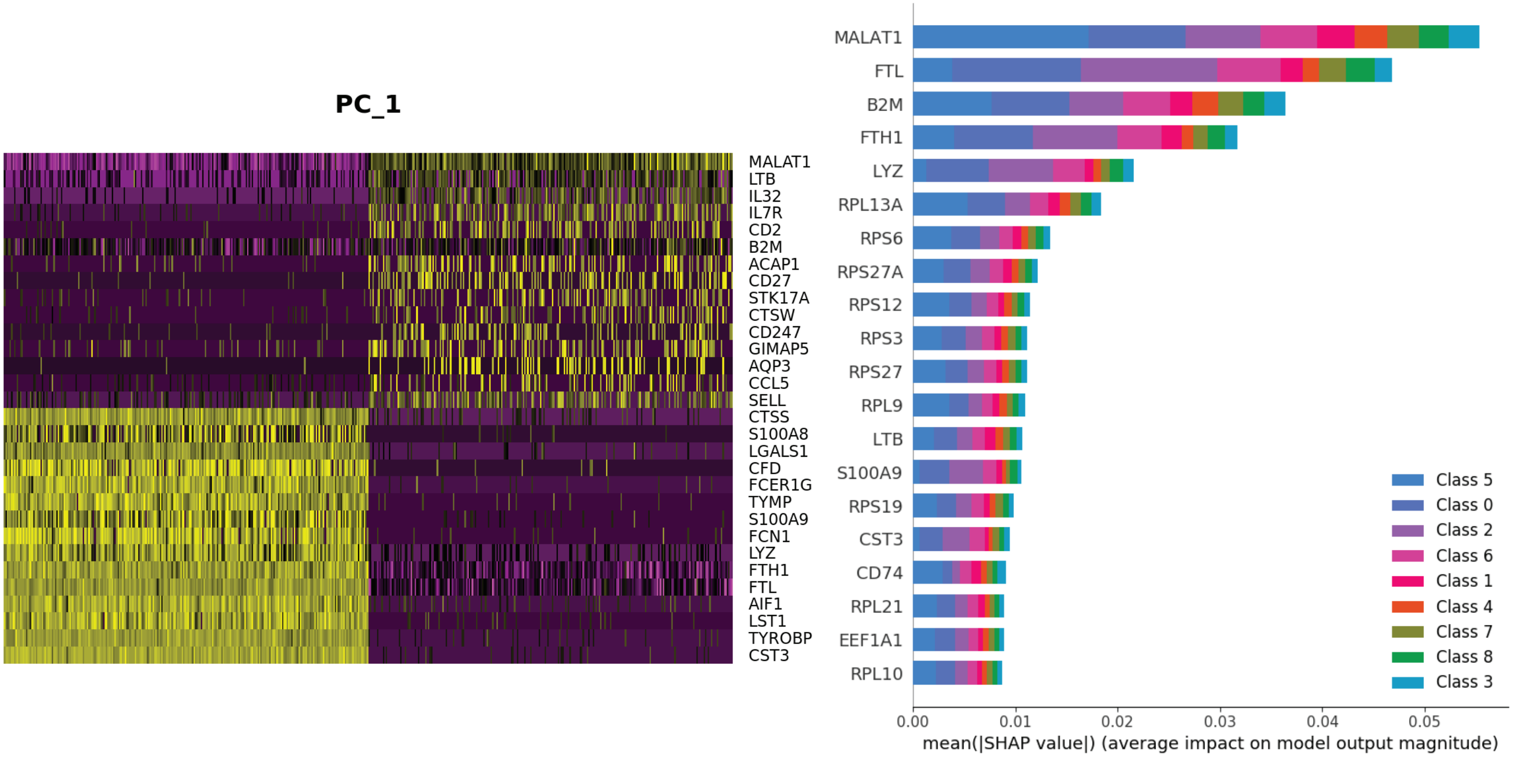
